# Risk of multiple primary cancer diagnosis over age in families of Li-Fraumeni syndrome: a single institution perspective

**DOI:** 10.1101/567693

**Authors:** Seung Jun Shin, Elissa B. Dodd, Fan Gao, Jasmina Bojadzieva, Jingxiao Chen, Xianhua Kong, Chris Amos, Jing Ning, Louise C. Strong, Wenyi Wang

## Abstract

Li-Fraumeni syndrome (LFS) is a rare autosomal dominant disorder associated with *TP53* germline mutations and an increased lifetime risk of multiple primary cancers (MPC). Penetrance estimation of time to first and second primary cancer within LFS remains challenging due to limited data availability and the difficulty of accounting for the characteristic effects of a primary cancer on the penetrance of a second primary cancer. Using a recurrent events survival modeling approach, we estimated penetrance for both first and second primary cancer diagnosis from a pediatric sarcoma cohort (number families=189; Single primary cancer, or SPC=771; MPC=87). We then validated the risk prediction performance using an independent clinical cohort of *TP53* tested individuals from MD Anderson (SPC=102; MPC=58). Our penetrance estimates are dependent on *TP53* genotype, sex, and, importantly, age of diagnosis (AoD) for the first PC. We observed the later the AoD is, the shorter the gap time is for this person to be diagnosed with a second PC. We achieved an Area under the ROC curve (AUC) of 0.77 for predicting individual outcomes of MPC vs. SPC. In summary, we have provided the first set of penetrance estimates for SPC and MPC for *TP53* mutation carriers, and demonstrated its accuracy for cancer risk assessment. Our open-source R package, LFSPRO, provides future risk estimates for SPC and MPC among *TP53* germline mutation carriers.

**Significance:** Our tool can be used to support clinical counseling of LFS cancer survivors for better health management.

## Introduction

Li-Fraumeni syndrome (LFS) is a familial cancer syndrome associated with germline *TP53* mutations and predisposing to a high lifetime probability of developing a wide spectrum of cancers^1^. Clinical management of families affected with *TP53* mutations remains a significant challenge. With the increasing success of treating cancer patients, many *TP53* mutation carriers survive their first primary cancers only to develop additional primary cancers throughout their lives. It has been estimated that the risk for second primary diagnosis can be as high as 50% for germline *TP53* mutation carriers^2^ and multiple malignancies have been previously observed in 43% of *TP53* carriers^3^. Many of these patients are aware that they are at increased risk of a second primary diagnosis, however the age of onset for a second primary is yet un-characterized in a statistically rigorous manner. Use of age-specific penetrance estimates of *TP53* mutation carriers may have implications for surveillance and clinical management. A rigorous cancer surveillance program for LFS at the University of Texas MD Anderson Cancer Center (MDACC) currently follows advanced screening protocols^4^. Counseling patients and family members could be further enhanced with a comprehensive cancer risk prediction tool that would enable the patients to understand or quantify their risk, lower anxiety for the unknown and improve adherence to screening.

Penetrance refers to the proportion of individuals carrying a deleterious variant of a disease predisposing gene (genotype) that also express clinical symptoms (phenotype) by a certain age. Penetrance can be incomplete and age-related, and precisely modeling it is of great importance to personalized risk assessment in medical genetics. Penetrance is further influenced by characteristics of the genotype, such as the specific impact that a variant has on a gene function and modifier factors^5–8^. Additionally, age-of-onset penetrance estimation is also important for characterizing genetic effects of a disease. The goal of this paper is to present a penetrance, estimated with information from all individuals from genetic pedigrees, and to independently validate risk for both first and second primary cancer diagnosis.

## Methods

### Model development

We defined the penetrance of primary cancer as the cumulative probability of developing the next primary cancer by a certain age given the mutation status of disease susceptibility variant and prior cancer history (e.g., previous primary cancer occurrence and diagnosis age). In this study, we estimated the penetrance specific to the first and the second primary cancer. The model for penetrance estimation is built based on survival modeling of recurrent events and on Mendelian inheritance property of genotypes, which allows us to model data from pedigree studies. The underlying theory for the statistical model has been previously developed^9^. In brief, we considered the multiple primary cancer occurrence in a randomly selected individual as a non-homogenous Poisson process (NHPP) and built the model with the following two major components: 1. Recurrent events modeling, which was devised to estimate the time varying hazard that fully characterizes the primary cancer occurrence process. We used a proportional hazard function where the baseline is a function of current age and the exponential component can incorporate covariates of interest. The model can thus consider effects from current age, cancer history or genetic factors when estimating the risk for next primary cancer development.

For the LFS data, we incorporate a covariate **X**(*t*) = {*G, S, G* × *S, D*(*t*), *G* × *D*(*t*)}^*T*^into the NHPP model, where *D*(*t*) is a time-dependent, but periodically fixed MPC variable that is coded as *t* > *T*_1_ and 0 otherwise. We propose the following multiplicative model for the conditional intensity function given **X**(*t*) as

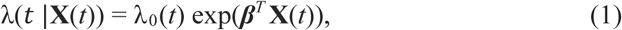

where ***β*** denotes the coefficient parameter that controls effects of covariate **X**(*t*) on the intensity and λ_0_(*t*) is a baseline intensity function. And 2. unknown genotypes among family members were imputed via the Elston-Stewart algorithm^10^, which significantly increases the statistical power for parameter estimation using all available cancer outcomes in these families. This approach improves computational efficiency by exploiting the Mendelian inheritance property when inferring missing genotypes in pedigrees. We also corrected for ascertainment bias due to selection of families though cancer-affected individuals and finally made inference on model parameters via a Markov chain Monte Carlo method. All 95% confidence intervals are 95% confidence bands derived from posterior distributions.

We specified the model with three main effects (*TP53* genotype (0 for non-carrier and 1 for carrier), gender (0 for female and 1 for male) and current cancer status (0 for no cancer diagnosis and 1 for one primary cancer diagnosis) and their interactions. We then computed the deviance information criterion (DIC) to identify the best set of covariates. We compare five different combinations of *G, S* and *D*(*t*). We observe that the simplest model with {*G, S, D*(*t*)} achieves the minimum DIC value. However, we decided to select the second best model in terms of the DIC, with {*G, S, G* × *S, D*(*t*), *G* × *D*(*t*)} as our final model since it has been reported that cancer status has different effects on cancer risk for mutation carriers and non-carriers^2^,^11^. Our model assumes similar penetrance of all cancer types due to limited number of patients for estimating penetrance for each cancer type separately. We considered death from any other cause or last follow-up as censoring events when estimating penetrance from cancer-free survival.

### Model training and validation study population

We used a MDACC pediatric sarcoma cohort data to train the model^12,13^. The data were collected based on probands with sarcoma diagnosed before age 16 and with at least 3 years after-diagnosis survival. The data collection was extended to the probands’ blood relatives, which includes the probands’ grandparents, parents, parental siblings, siblings and offspring and pedigrees could be further extended through affected relatives using a sequential sampling scheme^14^. For each individual, the gender and the diagnoses of any malignant cancer except the non-melanoma skin cancer were recorded from the date of birth until the date of death or last contact date. Medical records and death certificates confirmed cancer diagnoses, where possible. The primary cancer diagnoses were determined based on the histology and site information recorded for each cancer event. Mutation carrier status in this study was defined by PCR screening of exons 2-11 of the *TP53* gene from peripheral-blood cell samples. More information about mutation testing can be found elsewhere^15^. The final data was comprised 189 families and a total of 3,706 individuals, with a total of 964 (26.0%) individuals genotyped for *TP53* mutations status (**Table 1**). Among 570 (15.4%) primary cancer patients identified, 52 (1.4%) patients developed multiple primary cancer during follow-up. In this data set we have approximately equal number of cancer patients or healthy individuals for each gender (**Table 1**).

**Table 1:** Number of primary cancer patients by the *TP53* mutation status and sex in the training dataset: the MDACC pediatric sarcoma cohort data. Abbreviations: SPC, single primary cancer patients; MPC, multiple primary cancer patients.

|  |  | Wildtype | Mutation | Unknown |
| --- | --- | --- | --- | --- |
| Healthy individuals | Male | 295 | 9 | 1276 |
|  | Female | 341 | 8 | 1207 |
| SPC | Male | 105 | 25 | 139 |
|  | Female | 121 | 23 | 105 |
| MPC | Male | 3 | 14 | 8 |
|  | Female | 3 | 17 | 7 |
| Total number of individuals |  | 868 | 96 | 2742 |

For model prediction performance validation, we used an independent MDACC data set of prospectively followed families that were selected based off of clinical LFS criteria^3,16,17^ that includes both *TP53* germline mutation carrier families and wildtype families. The number of primary cancers in this data is summarized in **Table 4**. We only used the individuals with available genotype information for validation (**Table 4**).

### Prevalence and de novo mutation rate

We assume the *TP53* mutation follows Hardy-Weinberg equilibrium. The mutation prevalence is set at 0.0006^18^. Correspondingly, the frequencies for the three genotypes (homozygous reference, heterozygous and homozygous variant) are 0.9988004, 0.00059964 and 3.6e-07, respectively. We used 0.00012 as a default value of *de novo* mutation rate when evaluating the familywise likelihood^19^.

### Validation study design

We evaluated the model prediction performance on primary cancer risk using the average annual risk computed with our *TP53* penetrance estimates. The risk was calculated as the cumulative probability of developing the next primary cancer divided by the follow-up time. The receiver operating characteristic (ROC) curve was used to evaluate the sensitivity and specificity of predicting a primary cancer incidence using the estimated risk probability at various cutoffs. For Kaplan-Meier (KM) method-based risk prediction, we obtained KM survival functions for the time from date of birth to first primary cancer. These survival probabilities were then converted to penetrance estimate to compute the average annual risk. We used Jackknife to compute the standard errors of prediction performances^20,21^. In brief, each subsample was generated by omitting the *i*th family and the average under the curve (*AUC*) was calculated for this subsample as previously described. The standard error (*se*) was calculated using the Jackknife technique,

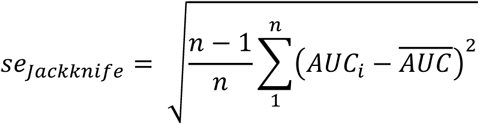

where *n* is the number of families, and 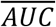 is the mean estimate of *AUC* values among all Jackknife subsamples.

## Results

### Age-of-onset penetrance curves for single- and multiple-primary cancers

**Table 1** provides a summary of the pediatric sarcoma cohort used to train our model. There are a total of 96 known *TP53* mutation carriers, among which 48 were diagnosed with one primary cancer and 31 were diagnosed with more than one primary cancer. There are 2,742 individuals who were not tested for *TP53* mutations, among which 244 were diagnosed with one primary cancer and 15 were diagnosed with more than one primary cancer. **Table 2** provides the coefficient estimates for all covariates including sex, genotype, cancer diagnosis and the interaction terms. As expected, the risk of the first and second primary cancers are strongly associated the *TP53* mutation status with a hazard ratio (HR) of *e*^*β*^= 27 (95% CI: 18, 40). Importantly, as illustrated in **Figure 1**, the *TP53* mutation carriers with a primary cancer diagnosis present a sharper increase in risk of having another primary cancer diagnosis over age than carriers who are still disease-free (HR=1.65, 95%CI: 1.10, 2.48). Such effect was not observed in the non-carriers (HR=0.82, 95%CI: 0.40, 1.48).

**Table 2:** Summary of covariate coefficients and their 95% confidence intervals estimated by our model.

| Covariate | Coefficient Estimate | 95% CI | Hazard Ratios |
| --- | --- | --- | --- |
| Genotype | 3.288 | (2.871, 3.687) | 26.782 |
| Sex | 0.027 | (-0.187, 0.241) | 1.027 |
| Genotype × Sex | -0.354 | (-0.817, 0.106) | 0.702 |
| Cancer status | -0.197 | (-0.929, 0.389) | 0.821 |
| Genotype × Cancer status | 0.700 | (-0.033, 1.548) | 2.014 |
| Cancer status + Genotype × Cancer status | 0.502 | (0.091, 0.908) | 1.652 |

**Figure 1:**
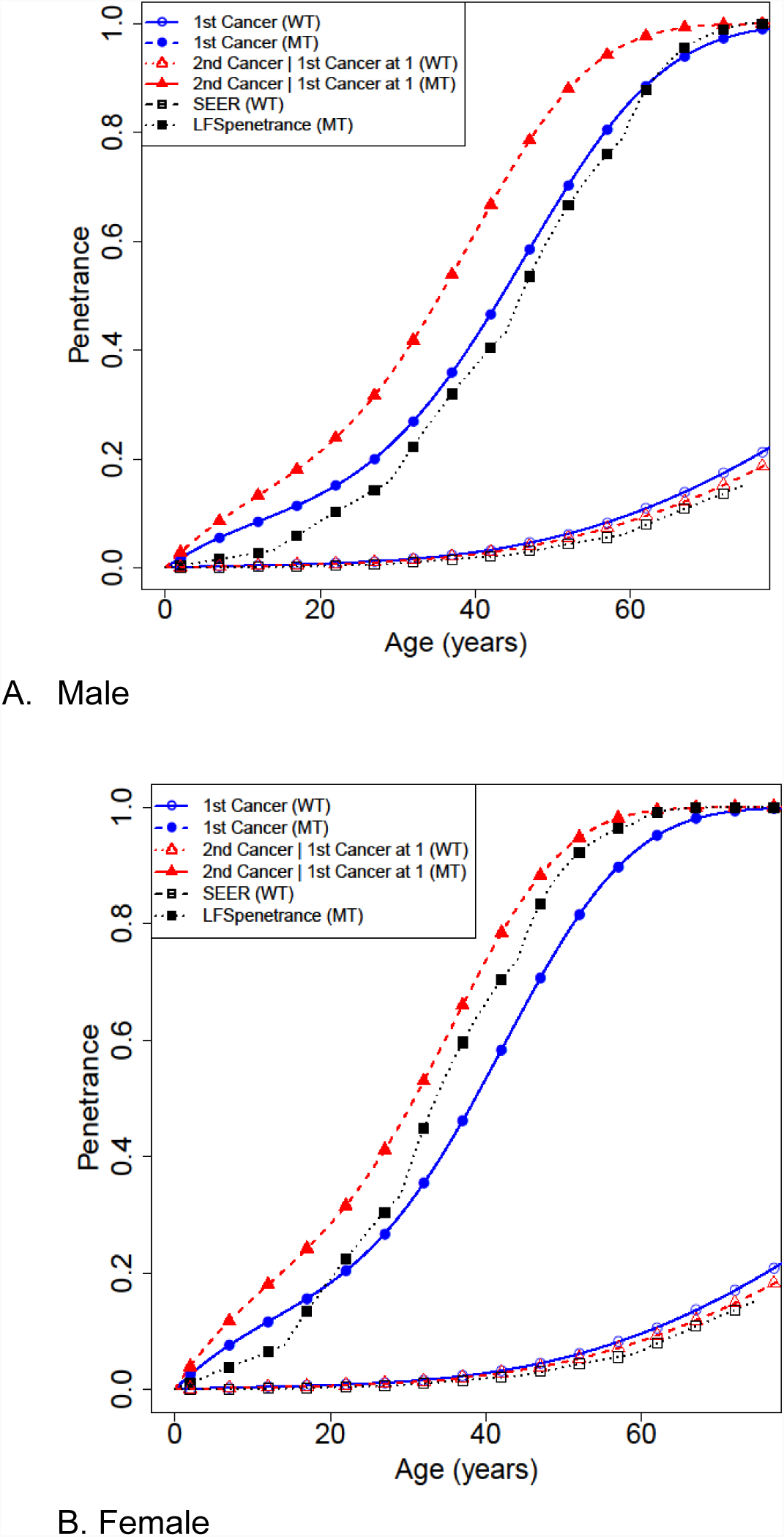
Penetrance estimates over time to the first cancer and time to the second cancer given the first cancer diagnosed at age 1, for A) male and B) female. Previously reported penetrance estimates for time to the first cancer are also shown for comparison: the SEER estimate for non-carriers, and the LFS penetrance for the first primary cancer for carriers by Wu et al., 2010^28^.

Using our model, we have obtained an accurate estimate of the onset of the first primary cancer by including cancer cases without genotype information from the family data. Among females, the HR for mutation carriers as compared to non-carriers is 26.8 (95%CI: 17.62, 39.88), while the HR among males for carriers versus non-carriers is 19.26 (95%CI: 13.14, 27.95). Consistent with previous results^15^, the number of mutation carriers are similar in males and females in this study (**Table 1**). However, the estimated cancer risks are different in early ages between males and females (cumulative risk by age 30 at 0.239 for male, and 0.317 for female), possibly due to the early onset of breast cancer in females (**Figure 1**).

We have further obtained a novel set of penetrance estimates for the age-of-onset for second primary cancers among individuals who have had been diagnosed with a primary cancer. As presented in our hazard model (**Table 2**), this set of penetrance is dependent on the age of diagnosis (AoD) of the first primary cancer (PC). To illustrate the dynamics with AoD, **Figure 2** shows the median age-of-onset risk for MPC within an interval of 20 for the age of diagnosis of the prior SPC: 0-20, 21-40 and 41-60. Here we observe a sharper increase in risk of developing a second primary cancer over age for individuals who have a later age-of-onset in their first primary cancer. Correspondingly, **Table 3** shows our estimated median (at 50% probability) time-to-a second cancer diagnosis for the 20-year age intervals, for both females and males, e.g., for a female carrier, the median times are 29 years for the early age group (0-20), 14 years for the middle age group (21-40) and 6 years for the late age group (41-60). Interestingly, similar observations can be made with non-carriers (**Figure 2**). Therefore our novel SPC/MPC penetrance estimates allows us to observe an age-dependent effect in the diagnosis of MPC in our cohort.

**Table 3:**
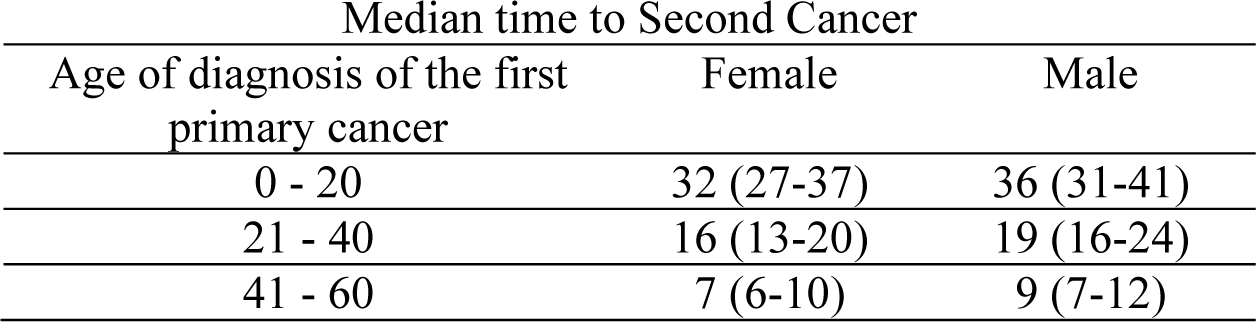
Median second primary cancer-free times (in years) since the first primary cancer diagnosis age and their 95% confidence intervals (in parenthesis) estimated for *TP53* mutation carriers, stratified by sex and age of diagnosis for the first primary cancer.

| Median time to Second Cancer |  |  |
| --- | --- | --- |
| Age of diagnosis of the first<br>primary cancer | Female | Male |
| 0 - 20 | 32 (27-37) | 36 (31-41) |
| 21 - 40 | 16 (13-20) | 19 (16-24) |
| 41 - 60 | 7 (6-10) | 9 (7-12) |

**Figure 2:**
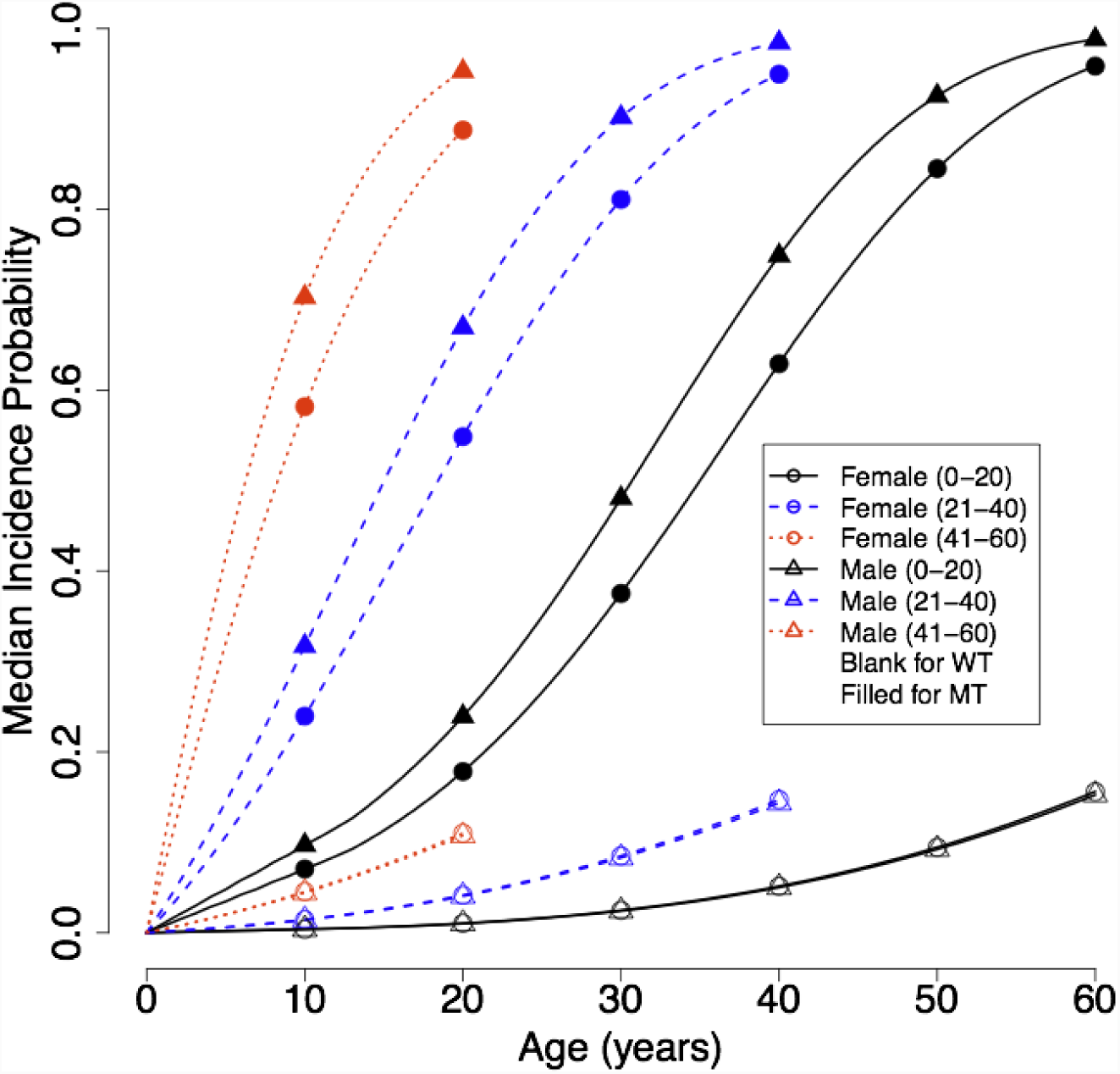
Illustration of the effect of second primary cancer on age-dependent penetrance estimates using median incidence probabilities in time windows of 20 years: 0-20, 21-40, 41-60. The x-axis denotes gap time, which is the number of years from the onset of the first primary cancer.

### Validation of risk prediction for first and second primary cancers

**Table 4** shows an overview of the validation population used for assessing the cancer risk prediction performance of the estimated penetrance. This dataset is not used for model development or model parameter estimation and hence serves as an independent test for risk prediction performance. We used individuals with known mutation status and cancer outcomes: 74 SPC and 55 MPC among the carriers, and 28 SPC and 3 MPC among the non-carriers. We evaluated the performance of our penetrance estimates by two types of risk assessment: 1). We evaluated the risk probability estimated for first primary cancer diagnosis; and 2) we evaluated the risk of second primary cancer diagnosis among individuals who have already had a first primary cancer diagnosis. As shown in **Figure 3**, our penetrance estimates achieved AUCs of 0.73 (standard error: 0.031) and 0.77 (standard error: 0.040) when predicting the first or the second primary cancer diagnosis, respectively. Our prediction for the first primary cancer diagnosis outperformed the commonly used KM method with a corresponding AUC of 0.67 (standard error: 0.036).

**Table 4:** Number of primary cancer patients by the *TP53* mutation status and Sex in the validation dataset: the MDACC prospective clinical cohort. Abbreviations: SPC, single primary cancer patients; MPC, multiple primary cancer patients.

|  |  | Wildtype | Mutation |
| --- | --- | --- | --- |
| Healthy individuals | Male | 114 | 40 |
|  | Female | 136 | 30 |
| SPC | Male | 16 | 31 |
|  | Female | 12 | 43 |
| MPC | Male | 1 | 21 |
|  | Female | 2 | 34 |
| Total number of individuals |  | 281 | 199 |

**Figure 3:**
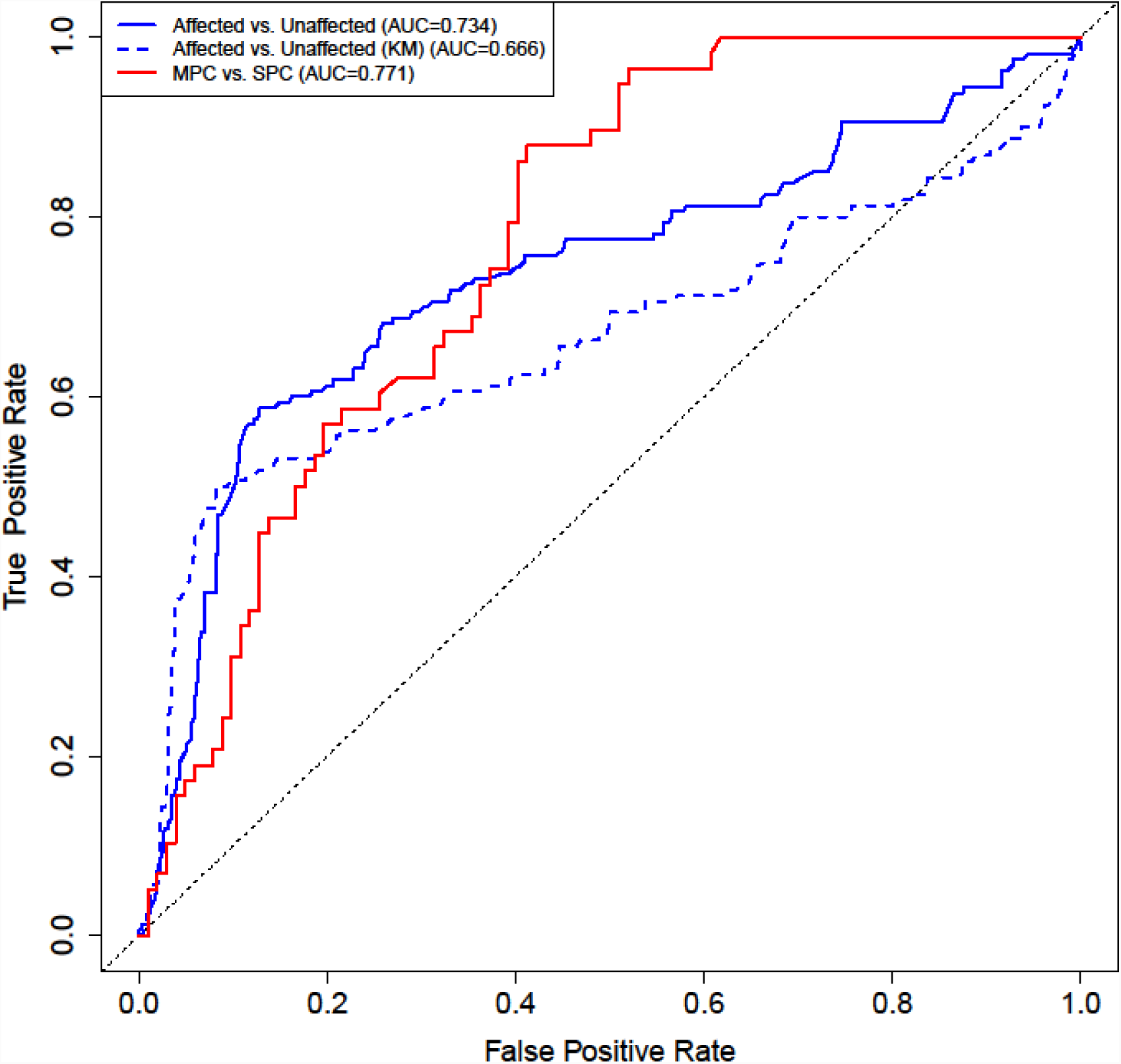
Comparison of validation performance between our multiple primary cancer-specific penetrance and those estimated from Kaplan-Meier (KM) method in predicting the first or the second primary cancer occurrence using the MDACC prospective data. Sample size: n(Affected)=160, n(Unaffected)=320, n(MPC)=58, n(SPC)=102.

## Discussion

This study presents a new set of penetrance for the first or second primary cancer diagnosis in patients with LFS and validated its risk prediction performance through an independent LFS dataset. In contrast with previous studies^2,15^, our NHPP model allowed us to utilize information from all family members, like sex, genotype if available, and age of diagnosis of the first primary cancer, by properly exploiting the family structure, using patient data with or without mutation test results. Including all individuals, regardless of testing, increased our training data sample size from 311 tested cancer patients to 570 total cancer patients, which substantially increased the statistical power for more accurate parameter estimation. Our final penetrance estimates for second diagnosis are age-of-diagnosis-dependent and varied with *TP53* genotype and sex. Based on the new penetrance estimates, we observed the risk of second cancer diagnosis increased with older age of first cancer diagnosis. The penetrance of our model demonstrated a better risk prediction performance as compared to classical nonparametric methods, such as KM, for survival outcomes, as shown in the validation results. We have integrated the new penetrance estimates in our risk prediction software LFSPRO as LFSPR_2.0.0 to provide risk estimates, which is freely available at https://bioinformatics.mdanderson.org/public-software/lfspro/.

Stringent surveillance recommendations for *TP53* mutation carriers have been established that includes annual whole body MRI (WB-MRI) and brain MRI, among other screening exams, for early detection of tumors^4^. Studies have shown that this intensive cancer surveillance protocol has led to the early detection of primary cancers usually only requiring resection instead of chemotherapy and/or radiation, both of which have potential for contributing to treatment related late effects^2^. Once identified early, treatment for carriers with a new primary can be quickly assessed increasing the likelihood of a positive outcome after early diagnosis for participants, which has been stated as a key benefit for continued screening^22,23^. However, clinical studies of this rigorous screening protocol have reported to have psychosocial drawbacks^22,23^. Nevertheless, early detection and peace of mind after results disclosure are noted as benefits gained through the screening process that outweigh the drawbacks^23^. Psychosocial studies assessed after long term participation in surveillance programs are not yet available to determine if burnout is an issue. Implementation of age-specific penetrance estimation in LFS screening programs could give genetic counselors and clinicians an opportunity to provide a more complete picture of predicted risk to time of first or secondary primary to their patients. Since secondary primaries are estimated to occur in 50% of carriers^2^, patients are encouraged to maintain the rigorous LFS screening protocol. The open-source R package, LFSPRO, which estimates the probability of an individual being a *TP53* germline mutation carrier, has been expanded to also estimate risk to either first or second primary cancer (lfspro.mode function with mode= “mpc”) within 5, 10, 15 and 20 years. We are currently acquiring feedback on LFSPRO’s utility within the MDACC Li-Fraumeni Education and Early Detection (LEAD) program^23,24^ which consists of a multidisciplinary team that works together to perform LFS screening protocols, analyze screening results and discuss future recommendations with the patients. Our goal is for LFSPRO to be used by genetic counselors and LFS clinicians as a tool to tailor their discussion of early cancer risks with their patients. However, at this time, our R package does not provide prediction beyond the second primary cancer or the recurrence of a primary cancer.

Our MPC penetrance estimation for the MDACC LFS cohorts is the first step in forming penetrance estimation within the established LFS community. Before becoming a clinical tool there are many factors that still need to be considered. First, we collapsed all cancer types into a single one and did not investigate the cause-specific penetrance for second primary cancer. Previous studies have shown the risk estimates varied among different cancer types, with breast cancer risk being dominant among female carriers^2,25^. Second, we did not incorporate the effect of treatment in our modeling because of limited treatment information in the pediatric sarcoma cohort data, which is a prevalent issue in most datasets collected to date. Previous studies^26,27^ have shown that cancer treatment may be a risk modifier for time to next cancer. Also, treatment of patients differs over the span of the cohort study due to technological innovation making it more difficult to capture in one study. In our NHPP model framework, treatment effects can be added when available. Though treatment effects are currently not estimated, they are implicitly accounted for in terms of risk prediction, with the other parameter estimates absorbing the effect of treatments. This explains the good performance of our current model as the independent validation set was also collected at MDACC. One potential drawback is a direct application of our penetrance to other study populations may not fit.

In summary, our study provides age-specific penetrance estimates for first or second primary cancer in individuals with LFS and has successfully validated its discrimination power between primary cancer patients and cancer-free individuals through another LFS data set. These estimates have the potential to provide a more accurate primary cancer risk assessment for patients with LFS, especially for cancer survivors who desire a better risk management of any future cancer development.

## Acknowledgments

We thank Dr. Jialu Li for his contribution to this project.

## Author contributions

Concept and design: S.J. Shin, J.Ning, W. Wang

Development of methodology: S.J. Shin, J.Ning, W. Wang

Acquisition of data: J. Bojadzieva, X. Kong, L.C. Strong

Analysis and interpretation of data: S.J. Shin, E.B. Dodd, Fan Gao, Jingxiao Chen, Chris Amos, W. Wang

Writing, review and/or revision of the manuscript: S.J. Shin, E.B. Dodd, J. Bojadzieva, C. Amos, L.C. Strong, W. Wang

Administrative, technical, or material support: E. B. Dodd

Study Supervision: W. Wang

